# scRNAseq_KNIME workflow: A Customizable, Locally Executable, Interactive and Automated KNIME workflow for single-cell RNA seq

**DOI:** 10.1101/2023.01.14.524084

**Authors:** Samina Kausar, Muhammad Asif, Anaïs Baudot

## Abstract

**Summary:** Single-cell RNA sequencing (scRNA-seq) is nowadays widely used to measure gene expression in individual cells, but meaningful biological interpretation of the generated scRNA-seq data remains a complicated task. Indeed, expertise in both the biological domain under study, statistics, and computer programming are prerequisite for thorough analysis of scRNA-seq data. However, biological experts may lack data science expertise, and bioinformatician’s limited understanding of the biology may lead to time-consuming iterations.

A user-friendly and automated workflow with possibility for customization is hence of a wide interest for both the biological and bioinformatics communities, and for their fruitful collaborations. Here, we propose a locally installable, user-friendly, interactive, and automated workflow that allows the users to perform the main steps of scRNA-seq data analysis. The interface is composed of graphical entities dedicated to specific and modifiable tasks. It can easily be used by biologists and can also serve as a customizable basis for bioinformaticians.

**Availability and implementation:** The workflow is developed in KNIME; its tasks were defined by R scripts using KNIME R nodes. The workflow is publicly available at https://github.com/Saminakausar/scRNAseq_KNIME.

## Introduction and motivation

Single-cell RNA sequencing (scRNA-seq) aims to quantify the gene expression of individual cells. Each run of scRNA-seq generates large amounts of noisy and sparse data, making rigorous data analysis and biological interpretation challenging.

The main steps of a scRNA-seq analysis are quality check, normalization, denoising and dimensionality reduction, clustering of cells, and identification of differentially expressed genes. All these steps are usually coupled with extensive data visualization. In bioinformatics, R and Python languages are frequently used to develop pipelines for the analysis of biological datasets. The Seurat [1] and Scanpy [2] packages, developed in R and Python, respectively, have been widely adopted for scRNA-seq data analysis. These packages perform the different tasks necessary for scRNA-seq data analysis. For example, they allow the users to normalize raw counts (gene/feature counts matrix), apply Principal Component Analysis (PCA) for dimension reduction and different clustering methods to identify clusters of cells. However, using such packages requires prior knowledge of R or Python. This prerequisite expertise in computer programming hampers the engagement of biological experts in scRNA-seq data analysis, which limits reliable and robust biological interpretations.

In this context, several web tools have been developed [3–7]. These tools are accessible but present some limitations. First, some tools are restricted to few steps of the scRNA-seq analysis pipeline. For example, visnormsc performs only the normalization [8]. Second, web tools follow a strict and previously-defined architecture, thus preventing the users with expertise in computer programming customizing or extending the tool. Moreover, web tools process and analyze data on their hosted servers and often lack well-defined policy for data protection and integrity, which raises concerns related to data security. In addition, scRNA-seq dataset are large and can crash web applications, forcing the users to re-run the analyses from the beginning.

Frameworks such as snakemake [9], nextflow (https://www.nextflow.io/index.html), Galaxy (https://usegalaxy.org/), and Konstanz Information Miner (KNIME) [10] can also be employed to implement workflows composed of several data analysis steps. For example, WASP is a snakemake workflow dedicated to scRNA-seq data analysis [11]. Importantly, the installation, execution, and modification of such frameworks require programming expertise. KNIME is a free and open-source data analysis framework with graphical interface. KNIME allows the creation of locally installable and reusable workflows for big data analysis and visualization. For instance, Kausar et al. proposed a KNIME workflow for machine learning in drug design [12]. Here, we propose scRNAseq_KNIME workflow, a KNIME-based workflow encompassing the main steps of scRNAseq analysis. scRNAseq_KNIME workflow is interactive and automated. It is also executable locally, thus eliminating the concern of data security and integrity. Overall, scRNAseq_KNIME workflow can be used by researchers lacking programming skills to run a complete scRNAseq analysis. It can simultaneously be exploited by bioinformaticians as a basis to customize and extend each step of scRNA-seq data analysis.

### Implementation and scRNAseq_KNIME workflow architecture

scRNAseq_KNIME workflow is developed in KNIME. In KNIME, a specific task, for example reading/writing files, is defined by introducing a node. A node is both a graphical entity and a basic processing unit. Some nodes are associated with predefined tasks, for example a node dedicated to reading CSV files. Other nodes are associated with tasks that can be defined by writing scripts in different languages, such as R, Groovy, Matlab and Python.

The scRNAseq_KNIME workflow is implemented with a modular structure (Supplementary figure 1), in which each module is composed of KNIME predefined nodes and nodes defined by custom R scripts (Figure 1A). The nodes are connected to each other through edges defining their relationships. The input module takes 10x genomics files (barcode, genes/features and matrix files) as input. The Quality Control (QC) module then assesses the quality of input data and applies user-defined cutoffs to exclude low quality data. The data normalization and denoising module prepares data for further analysis and identifies highly variable features. The dimension reduction and clustering module allow users to apply linear and non-linear dimension reduction approaches and identify clusters of cells. In the next step, various statistical tests can be applied on the identified clusters to highlight important genes. The next module allows marker-based cell type annotation of the clusters. Interactive visualizations are provided by the final visualization module.

**Figure 1:**
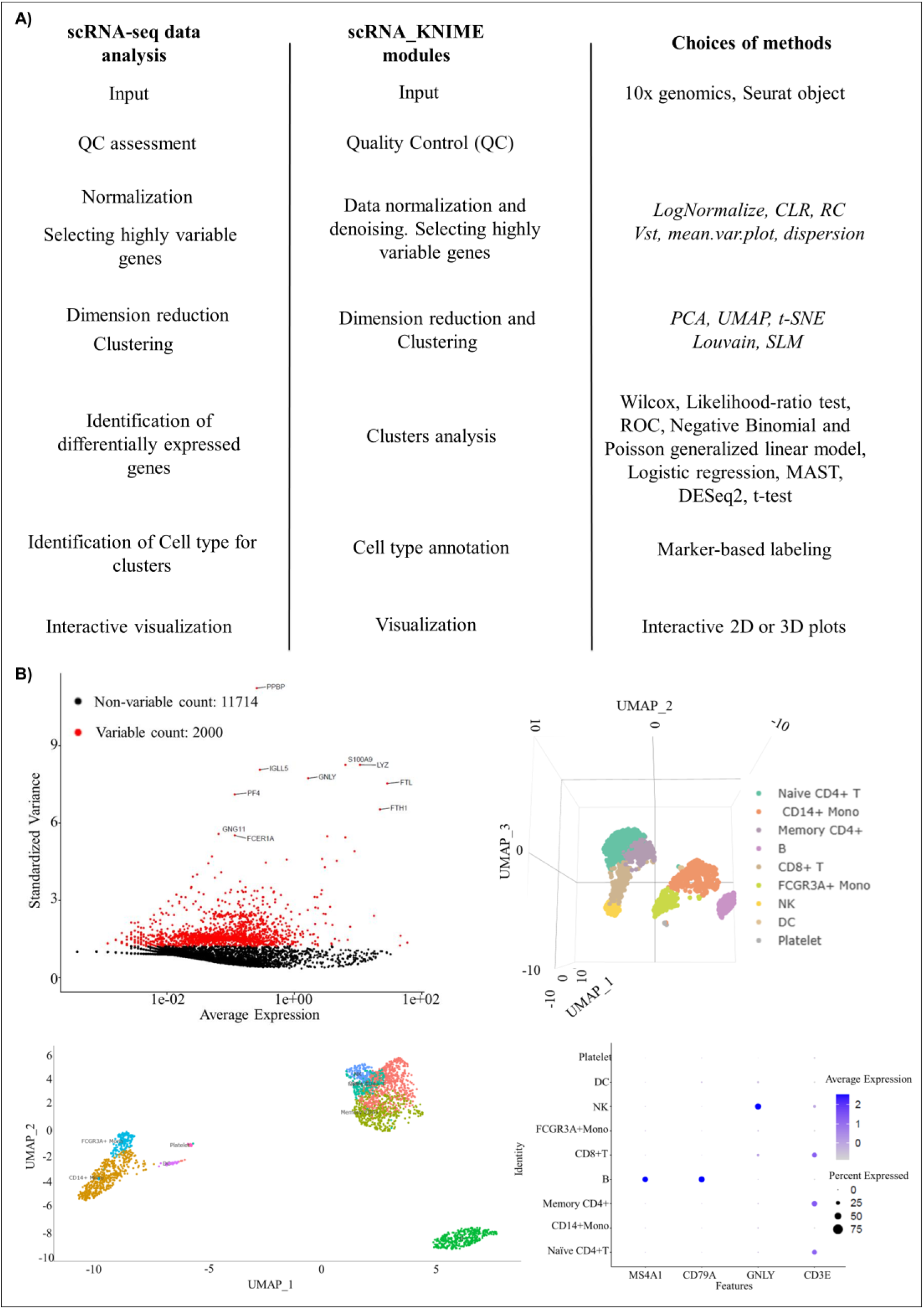
A) scRNAseq_KNIME workflow vs main steps of scRNA-seq data analysis workflow. MDS: Multidimensional scaling. B) Examples of interactive visualizations generated by the scRNAseq_KNIME workflow

scRNAseq_KNIME workflow provides extensive and interactive visualization. For example, clusters of cells can be visualized in 2D and 3D interactive plots. Many other data visualization options including VlnPlot, FeaturePlot, DotPlot, Heatmap, and interactive FeaturePlot are provided. Results of downstream analyses such as identification of differentially expressed genes are also provided through interactive data tables. To showcase the scRNAseq_KNIME workflow and its usage, we applied it to the analysis of the single-cell RNAseq dataset of Peripheral Blood Mononuclear Cells (PBMC), downloaded from 10x Genomics (Figure 1 B).

Overall, the scRNAseq_KNIME workflow implements the main steps required for scRNA-seq data analysis. In addition, each node of scRNAseq_KNIME workflow is interactive, and the users can test multiple parameters (Figure 1A and supplementary figure 2). Importantly, the R nodes can further be customized by modifying the R scripts.

The main features of scRNAseq_KNIME workflow are:

- Easy installation and execution: The workflow is developed with the KNIME platform, which comes with extensive documentation for its installation and execution. We further provide detailed documentation for running scRNAseq_KNIME workflow on Mac, Linux and Windows operating systems.
- Completeness: scRNAseq_KNIME workflow covers the main steps of scRNA-seq data analysis, allowing users (biologists and bioinformaticians) to run complete scRNA-seq data analysis and generate publication-ready plots.
- Possibility of customization: The scRNAseq_KNIME workflow modules are composed of R nodes containing R scripts. Users with expertise in R can customize R nodes by modifying the existing code.
- Modularity: Like snakemake, scRNAseq_KNIME workflow modules are independent from each other, meaning a change in one module of workflow does not require processing of previous modules.

scRNAseq_KNIME workflow is a locally installable, executable, and user-friendly framework that allows biologists to perform scRNA data analysis without knowing computer programming and statistical methods. The main steps of scRNA data analysis in scRNAseq_KNIME workflow can be customized by bioinformaticians by altering the code in R nodes of workflow.

