## Supplementary figures and images for "scRNAseq_KNIME workflow: A Customizable, Locally Executable, Interactive and Automated KNIME workflow for single-cell RNA seq"

### supplementary figure 1

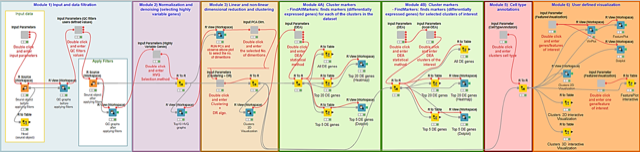
