## supplementary figure 2 for "scRNAseq_KNIME workflow: A Customizable, Locally Executable, Interactive and Automated KNIME workflow for single-cell RNA seq"

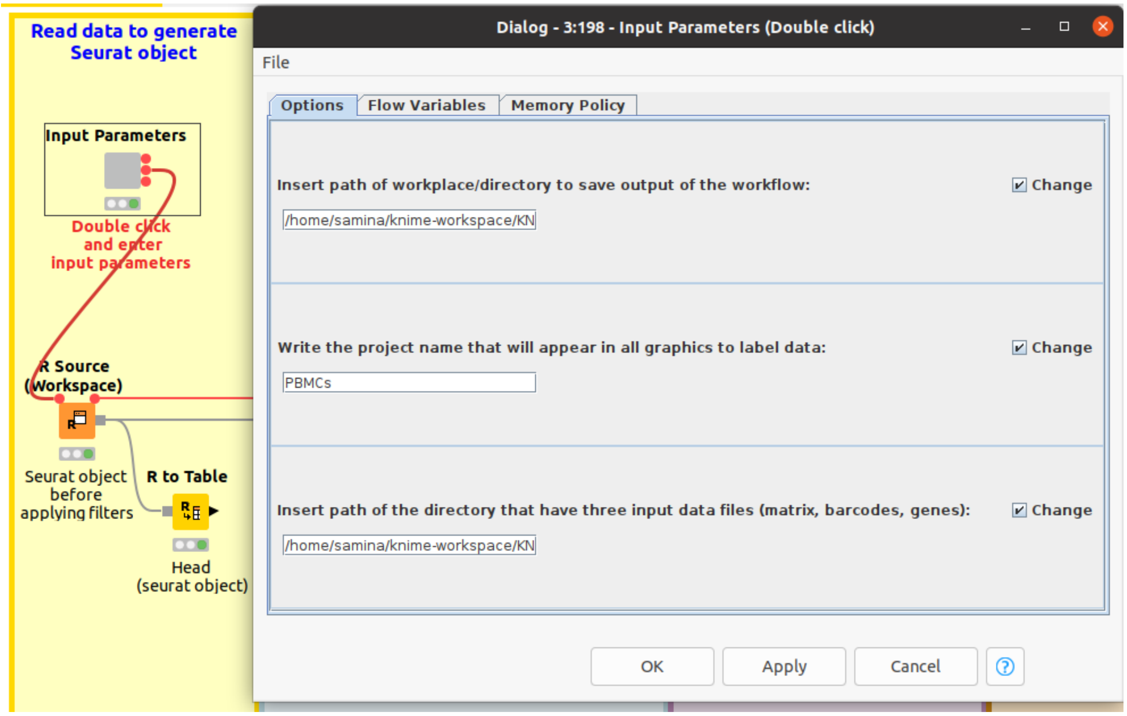


Supplementary figure 2: Example of input parameters configuration window, for the input data module
